# Optimizing the fitting initial condition for the parallel intrinsic diffusivity in NODDI: An extensive empirical evaluation

**DOI:** 10.1101/630541

**Authors:** Jose M. Guerrero, Nagesh Adluru, Barbara B. Bendlin, H. Hill Goldsmith, Stacey M. Schaefer, Richard J. Davidson, Steven R. Kecskemeti, Hui Zhang, Andrew L. Alexander

**Affiliations:** Department of Medical Physics, University of Wisconsin - Madison; Waisman Center, University of Wisconsin - Madison; Department of Medicine, University of Wisconsin - Madison; Department of Psychology, University of Wisconsin - Madison; Center for Healthy Minds, University of Wisconsin - Madison; Department of Computer Science, University College London

## Abstract

**Purpose:** NODDI is widely used in parameterizing microstructural brain properties. The model includes three signal compartments: intracellular, extracellular, and free water. The neurite compartment intrinsic parallel diffusivity (*d*_‖_) is set to 1.7 *µ*m^2^⋅ms^−1^, though the effects of this assumption have not been extensively explored. This work seeks to optimize *d*_‖_ by minimizing the model residuals.

**Methods:** The model residuals were evaluated in function of *d*_‖_ over the range from 0.5 to 3.0 *µ*m^2^⋅ms^−1^. This was done with respect to tissue type (i.e., white matter versus gray matter), sex, age (infancy to late adulthood), and diffusion-weighting protocol (maximum b-value). Variation in the estimated parameters with respect to *d*_‖_ was also explored.

**Results:** Results show the optimum *d*_‖_ is significantly lower for gray matter relative to 1.7 *µ*m^2^⋅ms^−1^ and to white matter. Infants showed significantly decreased optimum *d*_‖_ in gray and white matter. Minor optimum *d*_‖_ differences were observed versus diffusion protocol. No significant sex effects were observed. Additionally, changes in *d*_‖_ resulted in significant changes to the estimated NODDI parameters.

**Conclusion:** Future implementations of NODDI would benefit from *d*_‖_ optimization, particularly when investigating young populations and/or gray matter.

## Introduction

In diffusion weighted magnetic resonance imaging (dMRI), biophysical models are used for relating the dMRI signal to microstructural properties in white and gray matter [1–7]. Neurite orientation dispersion and density imaging (NODDI) [7], separates the brain tissue microstructure landscape into three compartments: intracellular space or neurites (axons, dendrites), extracellular tissue matrix, and a free water compartment. In spite of its shortcomings, much like the case of other techniques such as diffusion tensor imaging (DTI), NODDI offers useful information and has been widely used in the investigation of brain tissue microstructure as a function of early development, cognitive function and aging as well as a number of neurological conditions [8–13].

Biophysical modeling relies on simplifying assumptions about the tissue properties. Besides the separation of tissue into three compartments, the NODDI model is characterized by the following features or assumptions. Each compartment is represented by its own normalized signal and volume fraction. Water exchange between compartments is assumed negligible. Neurites are modeled as sticks (cylinders of zero radius) for capturing highly anisotropic architecture of neuronal tissue. Diffusion inside the neurites is described by a diffusivity parallel to the sticks, which is referred to as the *intrinsic diffusivity*, *d*_‖_, and zero diffusivity perpendicular to them. The orientation distribution function (ODF) of the sticks at each voxel is modeled by an axially symmetric Watson distribution, *W* [14], which itself is characterized by a concentration parameter *κ* and mean orientation ***µ***. Highly aligned sticks like those seen in white matter bundles are reflected by high *κ* values, while highly dispersed sticks like those seen in gray matter fibers are reflected by low *κ*. The extra-neurite compartment is directionally correlated with the neurite ODF, and modeled as a Gaussian anisotropic compartment.

The local parallel diffusivity of the extracellular space is set equal to the intra-neurite intrinsic diffusivity, *d*_‖_, whereas the perpendicular diffusivity *d*_⊥_ is related to the neurite water fraction, *f*_ic_, and *d*_‖_ by the mean-field tortuosity model [15] as *d*_⊥_ = (1 *− f*_ic_)*d*_‖_. The free-water compartment is modeled as having isotropic diffusion with free diffusivity *d*_iso_ = 3 *µ*m^2^⋅ms^−1^ and volume fraction *f*_iso_. The intrinsic diffusivity *d*_‖_ for NODDI is assumed to be 1.7 *µ*m^2^⋅ms^−1^. This is selected to be a biologically reasonable value, which approximates the mean parallel diffusivity from DTI in a healthy coherent white matter region [1]. The parameters that are estimated from acquired data using non-linear gradient descent and heuristic initializations are the water fraction of the neurite compartment (*f*_ic_), the concentration (*κ*) and mean orientation (***µ***) of the Watson distribution. The signal *S*(*b*, **g**) from the unit diffusion gradient direction **g** for sticks oriented along unit vector **n** and *b*-matrix (*b***gg**^*t*^) is given by

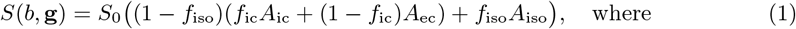

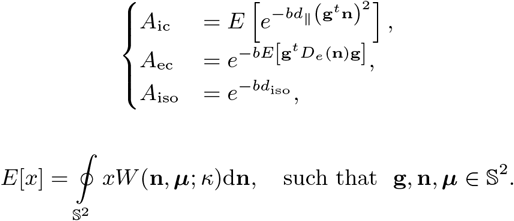

*A*_ic_, *A*_ec_, and *A*_iso_, are the intra-cellular, extra-cellular, and free-water isotropic compartments signal contributions respectively. *W* (**n***, **µ***; *κ*) is the Watson distribution with *κ* concentration and oriented along ***µ***. *S*_0_ is the un-attenuated signal i.e. *S*(0, 1), and *D*_*e*_(**n**) = *f*_ic_*d*_‖_ **nn**^*t*^ + (1 *− f*_ic_) *d*_‖_*I*_3_ is the axially symmetric extra-cellular apparent diffusion tensor.

Recently, the model assumptions have been a topic of discussion in the field. The more relevant discussions have focused around the fixed parallel intrinsic diffusivity and equality between parallel intrinsic diffusivity of the extra- and intra-cellular compartment [16]. Of the two, the equality assumption is the more difficult to assess, but has been explored in several reports. While no consensus has been reached, most reports suggest that the intra-cellular parallel intrinsic diffusivity is larger than the extra-cellular one [17–20]. Yet, this may depend on tissue type [21] and most studies have focused on white matter. Also, some sustain that the differences may not be substantial and independent validation experiments are needed [16].

With respect to the fixed diffusivity assumption, [22] proposed a framework for relaxing the fixed constraints. The study reported that microscopic parallel diffusivities varied across the brain, and that white matter values where considerably larger than that assumed by NODDI. It is important to note, however, that the ability to “estimate intrinsic diffusivity” in [22] comes at a cost, which is the reduction to two-compartment model. In this sense, then, the model in [22] is not fully comparable to the model in NODDI, since the former gives up on estimating the CSF volume fraction. Others [23, 24] have also relaxed the fixed diffusivity constraint and made it a free parameter. However, this resulted in unwanted effects on the other parameters in the form of unstable and degenerate estimates. Originally, it was considered unlikely that variation in *d*_‖_ across regions and subjects was significant enough to remove trends in the estimated parameters [1]. Additionally, the fixing of *d*_‖_ is necessary for stability in the parameter estimates and for speeding up convergence of the fitting procedure. Plus, the value that was chosen was the value that minimized the fitting errors for voxels in the midsagittal plane of the corpus callosum [1].

Taking into consideration the non-consensus on the equality assumption and the still widespread use of the technique, here we choose to build on earlier work [25] which investigated the assumption of fixed diffusivity. This consisted on the simple approach of optimizing the parallel intrinsic diffusivity based on the model residuals. Results suggested that the default value was reasonable in white matter, but it was sub-optimal in gray matter. While recent publications have found our method useful [26, 27], this earlier work only considered a single axial slice from three age matched participants and dMRI data acquired with the same imaging protocol. For this reason, we propose a more extensive implementation of our method for optimizing the intrinsic diffusivity that considers a diverse array of data in terms of age populations, imaging protocols, and is conducted across the full brain.

We hope for this extensive analysis to serve as a useful reference for the growing number of users of NODDI in making more conscious inferences based on the model parameter estimates.

## Materials and methods

### Data

Datasets acquired with multiple *b*-value sequences (suitable for implementing the NODDI technique [7]) were readily available for use in this work from a number of existing neuroimaging studies. These include imaging data from individuals with a broad range of ages and acquired with imaging protocols that vary in regards to number and magnitude of *b* values as well as number of diffusion encoding directions. dMRI sets include infants, adolescents, young adults, adults, and aging adults. All dMRI sets were collected on a 3T MR750 Discovery scanner (General Electric, Waukesha, WI). A brief description of each study is provided below and details are summarized in Table 1. All procedures for the included studies were approved by the University of Wisconsin - Madison Institutional Review Board.

**Table 1.** Relevant characteristics of studies from which data were used for this work.

| Study | Sex | Age | b-values [$\text{ms} \cdot \mu\text{m}^{-2}$] | Directions |
| --- | --- | --- | --- | --- |
| Neonates | 50 males 54 females | 1 month | 0.35, 0.8, 1.5 | 63 |
| Teen-I | 24 males 168 females | 11-15 years | 0.32, 0.8, 2.5 | 62 |
| Teen-II | 51 males 79 females | 14-20 years | 0.5, 0.8, 2.0 | 57 |
| Midlife-I | 57 males 89 females | 25-65 years | 0.5, 0.8, 2.0 | 57 |
| Midlife-II | 62 males 76 females | 25-74 years | 0.4, 1.2 | 76 |
| AD-risk | 18 males 53 females | 47-76 years | 0.3, 1.2, 2.7, 4.8, 7.5 | 105 |

#### Neonates study (Neonates)

Participants are from a study of neonatal white matter microstructure. Diffusion scans contain 6 non-diffusion weighted volumes and diffusion encoded along 63 directions. Other imaging parameters include: *TR/TE* = 8400*/*94ms, 2mm isotropic resolution.

#### Teen study (Teen-I)

Participants in this cohort were drawn from a study of emotion in adolescents. Diffusion scans contain 6 non-diffusion weighted volumes and diffusion encoded along 62 non-collinear directions. Other imaging parameters include: *TR/TE* = 8400*/*94 ms and 2 mm isotropic resolution.

#### Twin teen study (Teen-II)

Participants are from a cohort of 130 adolescent twins. Diffusion scans contain 6 non-diffusion weighted volumes and diffusion encoded along 57 directions. Other parameters include 2.0 mm isotropic resolution and *TR/T E* = 8000*/*66.2 ms.

#### Midlife meditation study (Midlife-I)

Participants in this cohort were drawn from a study of emotion regulation, asthma, and sleep part of the National Center for Complementary and Alternative Medicine (NCCAM). Diffusion scans contain 6 non-diffusion weighted volumes and diffusion encoded along 57 directions. Other parameters include 2.0 mm isotropic resolution and *TR/T E* = 8000*/*66.2 ms.

#### Midlife in the US study (Midlife-II)

Participants for this cohort were drawn from the Midlife in the United States (MIDUS), a national longitudinal study of health and well-being across the lifespan Refresher sample. Diffusion scans contain 4 non-diffusion weighted volumes and diffusion encoded along 76 non-collinear directions. Other imaging parameters include: *TR/TE* = 7000*/*69 ms and 2 mm isotropic resolution.

#### Preclinical Alzheimer’s disease risk study (AD-Risk)

Participants were cognitively unimpaired individuals with and without increased risk for Alzheimer’s disease recruited from the Wisconsin Registry for Alzheimer’s Prevention and Wisconsin Alzheimer’s Disease Research Center. Diffusion scans contain 7 non-diffusion weighted volumes and diffusion encoded along 105 non-collinear directions. Other imaging parameters include: *TR/TE* = 6500*/*102 ms, sagittal slices 3*mm* thick, and in-plane resolution of 2.5 mm *×* 2.5 mm.

### Intrinsic diffusivity optimization

*d*_‖_ was optimized by minimizing the model residuals. The search space was defined by the interval [0.5, 3.0] *µ*m^2^⋅ms^−1^ in increments of 0.1 *µ*m^2^⋅ms^−1^. For each of the 26 values, the model was fitted to the measured dMRI signal voxel by voxel using the Matlab (The MathWorks, Inc., Natick, MA) NODDI toolbox^1^. Predictions of the signal were then calculated at each voxel from the estimated parameters. With the measured and predicted signals for each *d*_‖_ setting, the root mean squared (RMS) residual was computed at each voxel. A linear search across the 26 different points was then performed for locating the value of *d*_‖_ corresponding to the lowest RMS residual value per voxel. The final result was a brain map of the *d*_‖_ that minimizes the RMS residual at each voxel (i.e. an optimized intrinsic diffusivity map).

### Tissue type segmentations

White matter (WM) and gray matter (GM) masks were obtained for each individual in order to probe the influence of tissue type on the fitting residuals. This was conducted by running FSL’s [28] FAST tool [29] with meand diffusivity (MD) and fractional anisotropy (FA) maps as input channels. FA and MD maps were obtained from tensor fits using a weighted least squares method. For the AD-risk study, the shells with b values of 4.8 and 7.5 ms*⋅µ*m^−2^ were excluded in the tensor fitting.

### Influences of age, sex, and protocol

The availability of data from the various studies allowed for selection of several subgroups that were organized according to age, sex, and protocol. With the data sets organized this way, the quality-of-fit analysis was performed for the following three cases:

#### Groups for age analysis

Subgroups of 16 participants (roughly half male and half female) were selected from three studies as follows: One group of 16 subjects age approximately one month from the Neonates study. One group of 16 subjects ages between 10 and 19 from the Teen-II study. Six groups, 16 subjects each, extracted from the Midlife-I study, for the six age categories of: 20-29, 30-39, 40-49, 50-59, and 60-65 years. Note that, except for the neonates, these data sets have matching protocols so that the main difference per category was age. In order to help disambiguate protocol from age influences, two additional scans were obtained for one adult: one with the infant protocol and one with the adult protocol.

#### Groups for sex analysis

From the Teen-I study, two subgroups one of 30 females and one of 30 males were selected. The two groups were matched by age (13 years old), so that the main difference between the groups was sex.

#### Groups for protocol analysis

Three groups of 16 subjects (roughly half females and half males) with ages ranging from 50-59 years were selected, one from the Midlife-I study, one from the Midlife-II study, and one from the AD-risk study. In this case, the assumed main difference between the groups was the acquisition protocol.

## Results

The results are organized as follows. **(1)** We first show how variation in *d*_‖_ translates to variability in the estimated parameters. **(2)** Then, the model RMS residuals, with respect to *d*_‖_ are shown to differ between tissue types. **(3)** This is followed by the presentation of voxel-wise optimized *d*_‖_ maps and the ways in which the optimum values are influenced by age, sex, protocol and tissue type. **(4)** Then, differences between optimized NODDI estimated parameter maps and those obtained with the default fixed *d*_‖_ are presented. Figures are best viewed in color.

### Estimated model parameters and *d*_‖_

Upon completion of the various model fits, the dependence of the estimated model parameters to variations in *d*_‖_ was explored. For all model parameter maps, mean values were calculated over WM and GM regions. Fig 1 shows these values plotted with respect to *d*_‖_. This analysis reveals a dependence on *d*_‖_ for all three parameters irrespective of the study as well as variation in the comparison of parameters among the studies. For example, for gray matter values of *d*_‖_ that are lower than the assumed value would weaken variation of the neurite density across the teen and adult subjects. On the other hand, lower values of *d*_‖_ in gray matter would enhance differences in the ODF concentration parameter across all studies.

**Fig 1.**
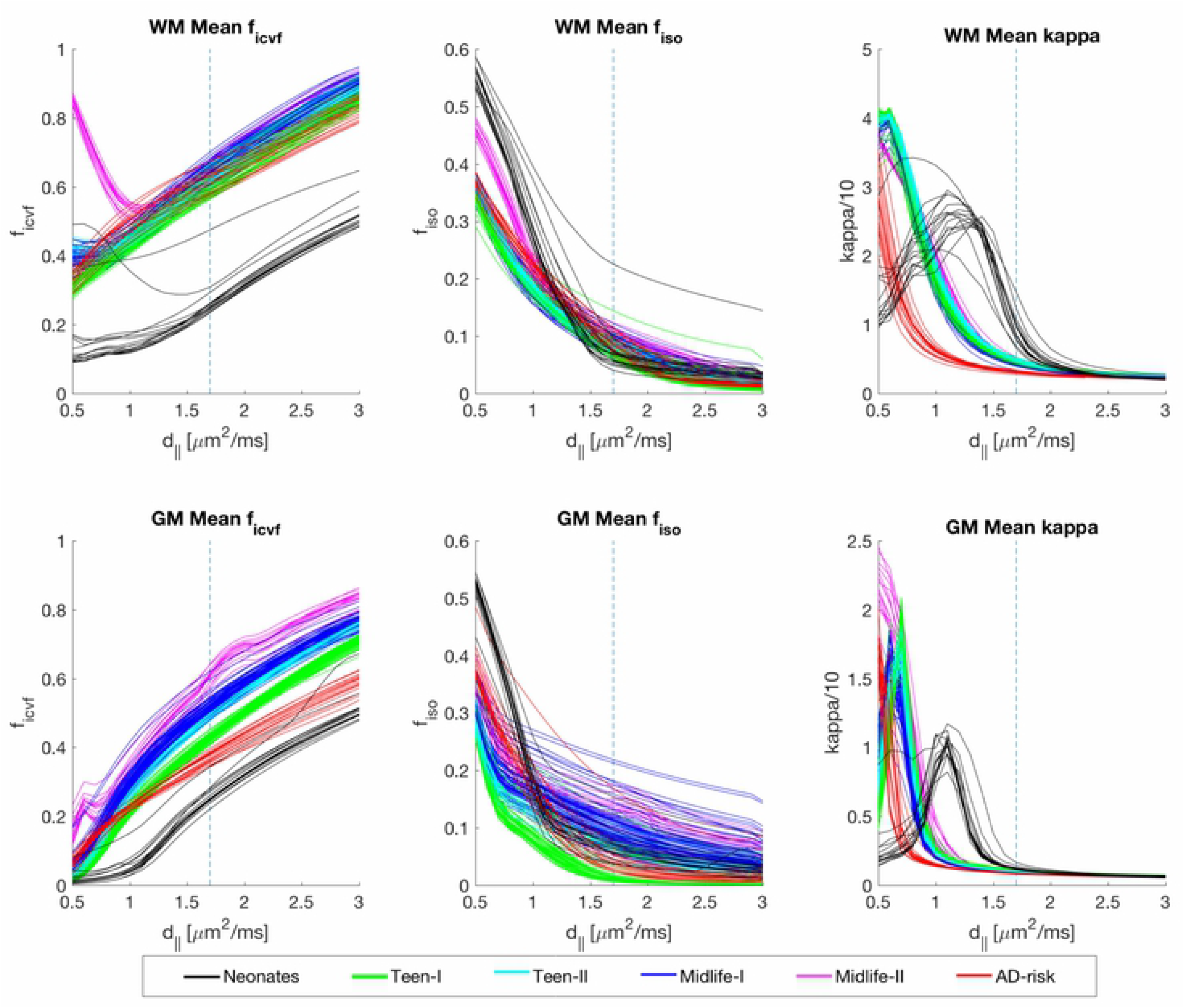
NODDI parameter trajectories with respect to *d*_‖_. For each parameter (Intra-cellular compartment volume fraction, *f*_ic_, isotropic compartment volume fraction, *f*_iso_, orientation concentration parameter, *κ*), the analysis is broken by white matter (WM) and gray matter (GM) regions. Each point on the curves represents the mean parameter over WM or GM at the specific *d* value. The default operating point is marked by the blue dashed vertical line.

### Model Residuals with respect to *d*_‖_

The values of *d*_‖_ that result in the closest agreement between the measured and predicted signals as dictated by the RMS residuals were explored next. For each of the resulting 26 RMS residual maps, mean values across WM and GM were calculated. These are plotted with respect to *d*_‖_ in Fig 2-A. These plots reveal that *d*_‖_ values in GM that achieve minimum RMS residuals deviate from the default setting (1.7 *µ*m^2^⋅ms^−1^) for all studies. For WM, the lowest values in the RMS residual curves occur in the neighborhood of the default setting. Notably, most WM curves, with the exception of the Neonate study, exhibit broad ranges of lowest values as compared to the majority of GM curves. The better defined minima in WM for the infants could be related to a maximum *b* value that better matches the characteristics of the young brain tissue (i.e. longer *T*_2_, low myelination, higher water content) such that diffusion weighting in the signal is more adequate. This is in line with the AD-risk study, which used a max *b*-value of 7.5 ms*⋅µ*m^−2^ and the WM RMS residual curves are noticeably more convex. The remaining studies have maximum *b* values that are likely on the low end of the optimal range for capturing effects of more restrictive intra-neurite environment, which could help explain the shallower curves in WM. The highest overall fitting errors occur for the Midlife-II and AD-Risk studies. These have the protocols that most deviate from the optimal NODDI protocol in terms of number and magnitude of b-values outlined in [7]. While for the Midlife-II scans the large fitting errors are directly related to a ‘bad’ model fit due to the low b values, the fitting errors for the AD-Risk scans may be more related to low signal to noise ratio in the images with the very large b values (4.8 and 7.5 ms*⋅µ*m^−2^).

**Fig 2.**
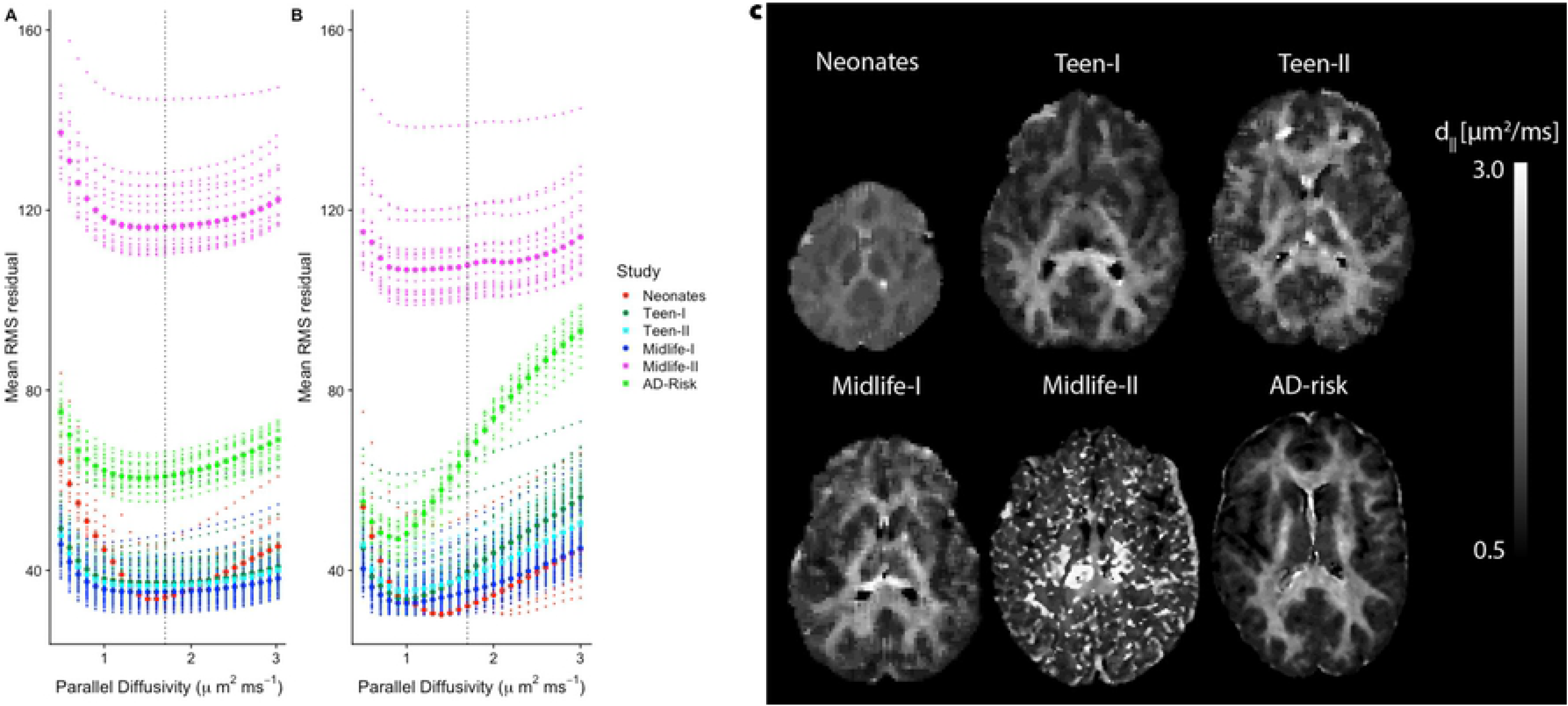
Model residuals with respect to *d*_‖_ and optimized *d*_‖_ maps. (**A, B**) Average root-mean-square (RMS) residual with respect to *d*_‖_ for all subjects in each study. Each of the small size dots represents the mean RMS residual over white matter (**A**) or gray matter(**B**) at the specific *d*_‖_ value. The large size dots represent the median value over all the subjects in the study at the specific *d*_‖_ value. The default operating point is marked by the blue dashed vertical line. (**C**) Axial view of optimized *d*_‖_ map for one subject selected from each of the studies.

#### Optimized *d*_‖_ maps

optimum intrinsic diffusivity whole brain maps were created by selecting at each voxel the value that corresponded to the smallest RMS residual. Resulting optimal *d*_‖_ maps were median filtered using a box kernel (size 3×3×3 in voxels). The filtering helps to enhance the underlying structure in the distribution of values between white and gray matter. The pattern is spatially consistent before filtering, but it is more difficult to appreciate due to the shallowness of the residual curves for white matter. Fig 2-B shows optimal *d*_‖_ maps for one subject selected randomly from each of the six studies. Except for the Midlife-II study, moderate to substantial contrast between WM and GM regions is apparent from these maps. The non-uniformly distributed *d*_‖_ in these maps suggests that a fixed diffusivity value may not be appropriate for all brain regions and all populations. Also, evident is the more noisy appearance of the map for the Midlife-II study participant, which could be explained by the low b value (see Table 1) and the shallow, unstable GM curves in Fig 2-A that correspond to this study. This is consistent with the more obvious WM-GM contrast in the map from the AD-risk study, which has the protocol with the highest b value.

#### Optimized *d*_‖_ and age

Optimal *d*_‖_ maps were computed for the cohort organized by age group. These maps were further masked into WM and GM regions and average optimal*d*_‖_ values were obtained for each region. Figure Fig 3-A shows the distributions of average optimal *d*_‖_ values according to age group. These plots show distinct distributions between WM and GM average optimal *d*_‖_ for all age groups greater than 10 years. The majority of WM optimal *d*_‖_ values are distributed around the default operating point (1.7 *µ*m^2^⋅ms^−1^), while all GM optimal *d*_‖_ values are reduced by at least 0.4 *µ*m^2^⋅ms^−1^. These trends are fairly consistent for all distributions corresponding to ages 10 years and above. For the group of less than 1 year (i.e. infants) there is a greater degree of closeness between the WM and GM distributions of average optimal *d*_‖_ in comparison to the rest of the age groups. In this case, optimal *d*_‖_ values fall approximately between 1.4 and 1.5 *µ*m^2^⋅ms^−1^ for WM and 1.2 and 1.3 *µ*m^2^⋅ms^−1^ for GM. For each age group, a pairwise t-test was conducted in order to assess statistical significance of the tissue-wise difference in average optimal *d*_‖_. The testing showed that for all groups the optimum *d*_‖_ for GM and WM were significantly different (*p* < 0.01). A multiple group test revealed that average optimal *d*_‖_ is significantly different between the infant and the rest of the older age groups in both WM and GM, while no significant differences were found between any of the other groups. The mean optimum *d*_‖_ values for the two additional scans on one adult, Fig 3-B, are in agreement with those values from same age group for both the infant and adult protocols, pointing to the fact that the observed trends are more a result of differences in age rather than in protocol.

**Fig 3.**
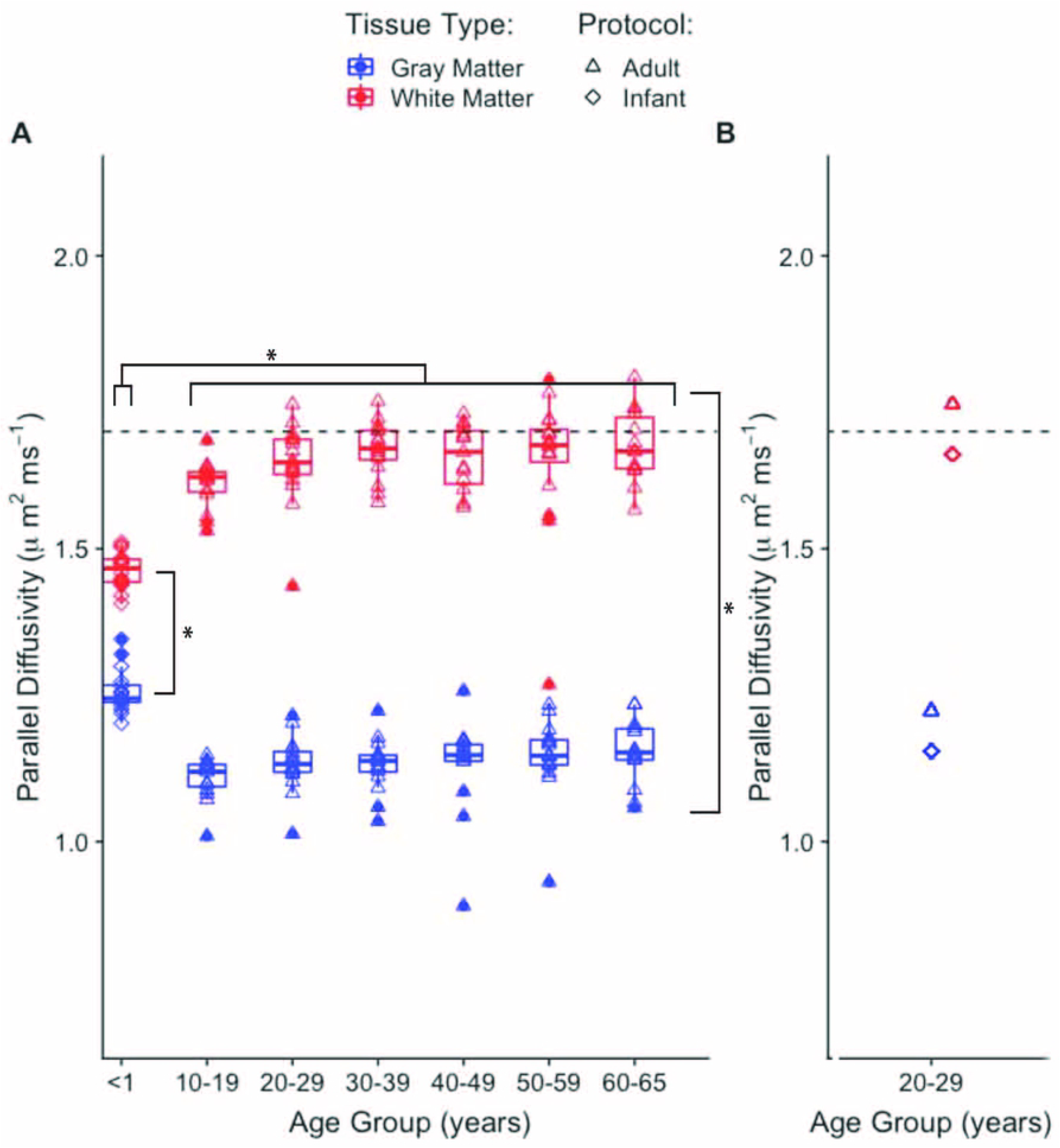
Optimized *d*_‖_ as function of age group and tissue type. (**A**) Mean value of optimal *d*_‖_ as function of age group and tissue type. The scanning protocol for the ¡1 year group is slightly different than that of the rest of the groups (Table 1). The numbers from scanning one adult with the two protocols are shown in **B**. The dashed horizontal line marks the default *d*_‖_ value.

#### Optimized *d*_‖_ and sex

Optimal *d*_‖_ maps were also computed for the cohort organized according to sex. Average optimal *d*_‖_ values were obtained across WM and GM regions. The distributions of average optimal *d*_‖_ values according to sex category revealed significantly different values between WM and GM with ranges that are consistent with the same age group (10-19 years) from the age-dependence analysis. Yet, no significant effects of sex were observed, a result that is compatible with the age-dependent analysis, which also showed no obvious split in optimal *d*_‖_ between the male and female participants.

#### Optimized *d*_‖_ and acquisition protocol

Finally, optimal *d*_‖_ maps were also computed for the cohort of subjects with data acquired under differing imaging protocols. Based on the observation that the age dependence analysis revealed no obvious age effects for ages 10 and above, data from the Teen-I study was also included in this cohort despite the unmatched age. This resulted in 4 protocol categories. Fig 4 shows the distribution of WM and GM average optimal *d*_‖_ values according to imaging protocol. Multiple group testing showed there exist significant differences between groups according to protocol in both white and gray matter. In WM, data sets from the protocol with the lowest b value present the lowest optimal *d*_‖_ when compared to the rest of the groups. However, this group also had the highest residuals in general in both WM and GM (see Fig 1 - Midlife-II study). The data sets from the groups with the highest b value protocol also show optimal *d*_‖_ values that are lower than the default operating point. In GM, this analysis reveals a seemingly decaying trend in optimal *d*_‖_ distributions with respect to maximum b value. Pair-wise t-tests revealed all distributions in GM are significantly shifted down compared to WM distributions, consistent with the observed trend in the previous age and sex comparisons.

**Fig 4.**
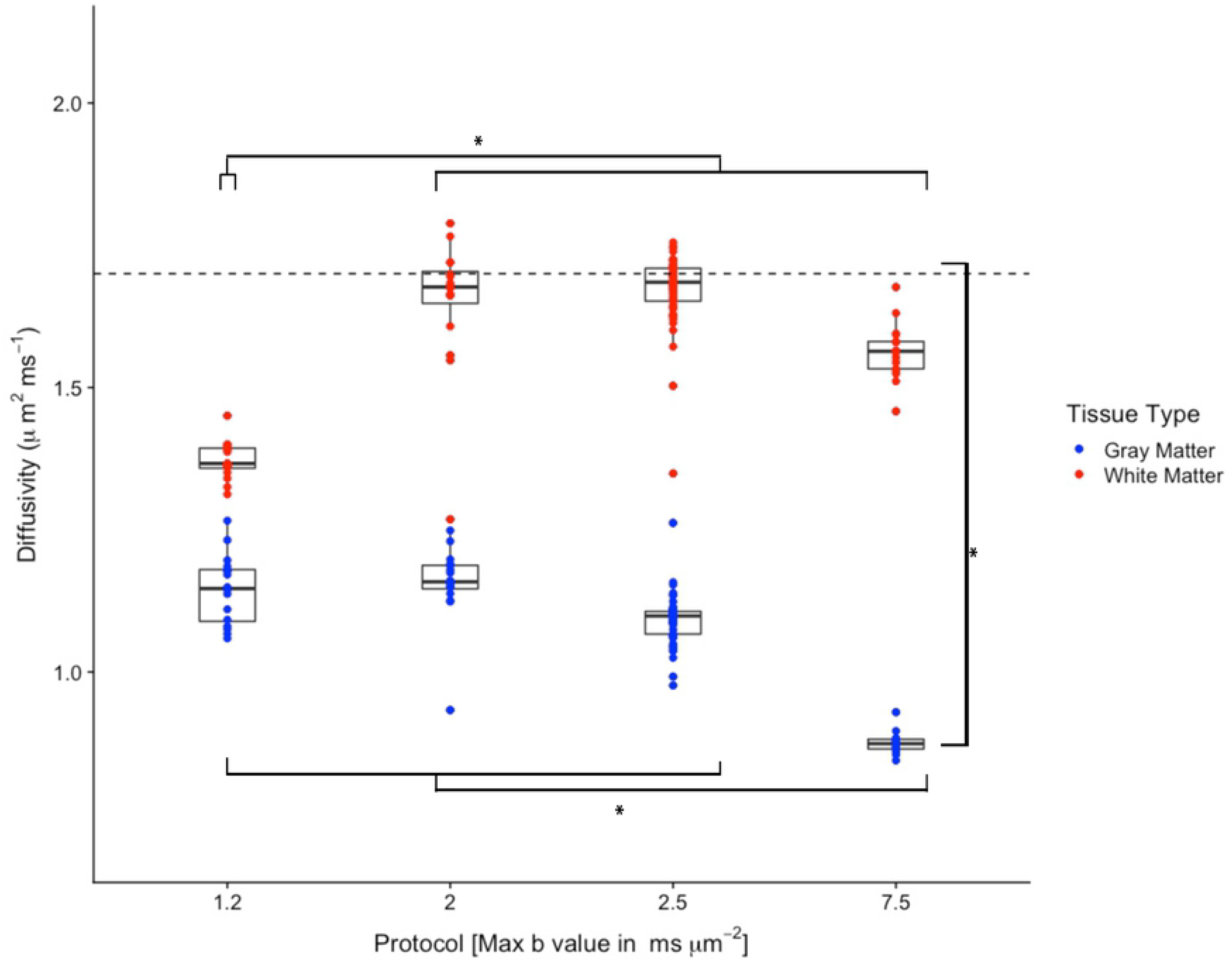
Optimized *d*_‖_ as function of imaging protocol. Mean value of optimal *d*_‖_ as function of imaging protocol and tissue type. The dashed horizontal line marks the default *d*_‖_ value.

#### Optimized NODDI parameter maps

Optimal NODDI parameter maps were estimated for all subjects by selecting the parameter value that corresponds to the optimal *d*_‖_ at each voxel. Fig 5 shows optimal *f*_icvf_, *f*_iso_, and *κ* maps for the same subjects in Fig 2-C. For comparison, Fig 5 also contains parameter maps obtained with the default setting for intrinsic diffusivity. Further, difference maps obtained by subtracting the default from the optimized maps are also shown. Mean difference results in white and gray matter for all subjects by study are shown on the right side column of Fig 5. Optimal NODDI does not seem to produce reasonable results for the Midlife-II study. This is likely a result of inadequate low b values used in the acquisition which are insufficient to capture the effects from the restricted space of the tissue and translate to a bad model fit in general. The difference maps point to existing biases, which are observed in the plots on the right. For instance, *f*_*icvf*_ mean differences in GM go from a couple percent to close to 30% of the range of possible values. *f*_*iso*_ mean differences in GM also reach values of higher than 20% of the possible values as shown by the 7.5 ms*⋅µ*m^−2^ b value subjects. And *κ* mean differences can be up to values of 1.6 in WM as shown by the infants group.

**Fig 5.**
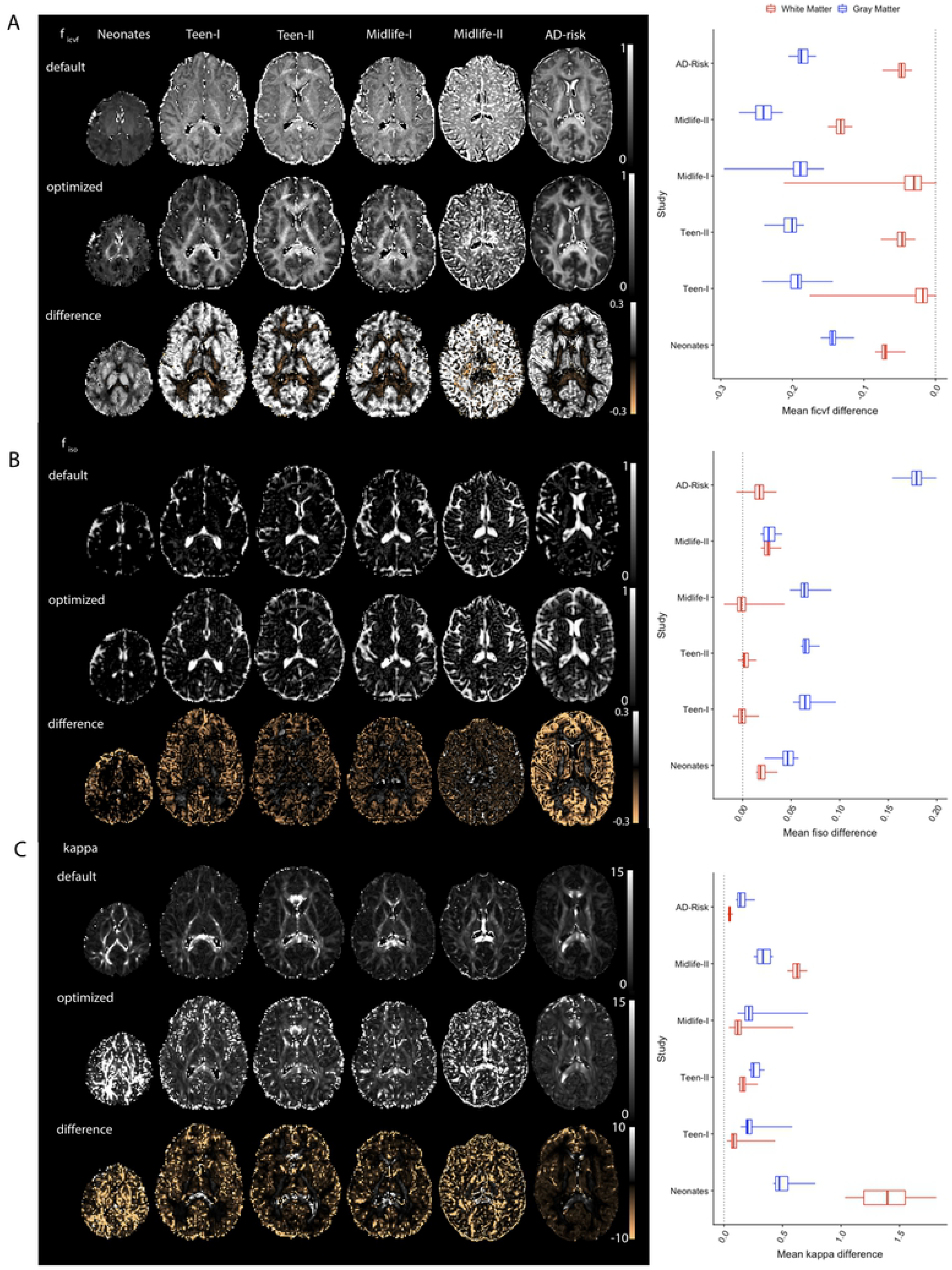
Optimized and default NODDI. Left column shows NODDI parameter maps from fits with the default *d*_‖_, with the optimized *d*_‖_, and optimized minus default signed difference from a subject in each study. Right column shows average difference over white and gray matter for all subjects in each of the studies. **(A)** Intra-cellular compartment volume fraction (*f*_*icvf*_), (**B**) Isotropic compartment volume fraction (*f*_*iso*_), (**B**) orientation concentration parameter (*κ*)

## Discussion

In this study we optimized the NODDI parallel intrinsic diffusivity (*d*_‖_) by minimizing the model residual for a diverse array of multi-shell dMRI data. The results suggests model assumptions for *d*_‖_ may be suboptimal for specific ages (i.e., infants) and also in gray matter. Although not examined, the optimal *d*_‖_ may also vary with pathology. We also observed that suboptimal *d*_‖_ leads to biases in the estimated NODDI parameters. Of particular interest is a drop of neurite density in gray matter observed in the optimized NODDI maps, a result that is consistent with findings in a recent study [30].

For gray matter, the optimal *d*_‖_ is significantly lower than 1.7 *µ*m^2^⋅ms^−1^. In white matter of the adult brain, values of the optimal *d*_‖_ hover around the default and below the range [1.9, 2.2] *µ*m^2^⋅ms^−1^ of intra-axonal diffusivities in white matter reported elsewhere [31], though, further analysis (see below) suggested high FA regions in the adult brain contained average optimal *d*_‖_ that falls in this range. It is important to note, however, that the ranges of residual minima in white matter were broad and shallow.

Further, a finer grain analysis indicates that protocol and age also have an impact on the optimal *d*_‖_, both in white and gray matter. The age-dependence analysis revealed that the newborn brain optimized *d*_‖_ in white and gray matter are closer in value compared to those in the adult brains. Both WM and GM values of optimized *d*_‖_ are different, however, from that used in recent studies [24, 32] that have implemented NODDI in the infant brain. The value in these studies was set to 2.0 *µ*m^2^⋅ms^−1^, likely because average DTI axial diffusivity in high FA regions (see below) of newborns is close to this number. Interestingly, at this setting, and using the 1.7 *µ*m^2^⋅ms^−1^ for the adult brain, nearly any difference between the infants ODF concentration parameter and that of the older age brains would be removed in gray matter. Using the optimal setting for *d*_‖_, would result in appreciable differences in ODF concentration parameter between the adults and the infants. On the other hand, using the optimal settings for *d*_‖_, would weaken the differences in intra-cellular volume fraction between the infant and the older subjects.

This analysis also showed that in the adult brain optimized intrinsic diffusivity values do not vary appreciably with age. However, optimum *d*_‖_ values in GM are much lower than those in WM and different from the default fixed value. With regards to imaging protocol, high b value and more diffusion weighted volumes appeared to yield less noisy and more stable optimal intrinsic diffusivity and NODDI parameter estimates.

In hindsight, the sub-optimality of the assumed *d*_‖_ in gray matter is not surprising since this value was originally estimated in the adult corpus callosum [1]. Also, sub-optimality of the current state of the model in gray matter might be related to the idea that the impermeable ‘stick’ representation of neurites is only adequate for myelinated axons but not for dendrites or non-myelinated axons, as others have suggested [33]. In general, however, the variation of optimal intrinsic diffusivity across tissue types is in agreement with findings of axial diffusivity variation across the brain reported in [30].

Studies have reported decreasing DTI axial diffusivity with age [34–36]. Thus, the trend of increasing optimum *d*_‖_ with age in WM seen in Fig 3-A was unsettling and prompted further investigation. For comparison, averages of DTI axial diffusivity over WM and GM were computed for all subjects in all age groups, Figure Fig 6-A,B. The resulting axial diffusivity age trajectories are in agreement with previous studies [34–36]. However, while these numbers pertain to the whole of white matter, regional differences in developmental trajectories of DTI quantities in the neonate brain have been observed [37]. In the infants, a further look into high FA (¿0.5) regions, which reduce to portions of the corpus callosum and the internal capsule, revealed that average optimal *d*_‖_ in these regions is comparable to that seen in the adult global WM. These regions in the infant are thought to be myelinated by one month after birth and to have higher fiber coherence than other white matter areas [37]. The lower FA regions (not shown) in the infant brain, which presumably reflect less or not-yet myelinated axons and or lower fiber coherence, exhibit values of average optimal *d*_‖_ that are similar to those of whole WM. For the older age groups, the axial diffusivity distributions in gray matter mimic those of the optimal *d*_‖_. For the infants, this is true for both the WM and GM distributions. Also, the optimal *d*_‖_ distribution comparison is less distinct for the infants than for the rest of the older age groups. Based on all this, it could be speculated that the neonatal gray matter neurites and white matter neurites are more similar than they are in the adults. Therefore, the model fit for less coherent, non-myelinated fibers in neonatal white matter would be more similar to the fit in the neonatal gray matter than to the fit in the adult whole WM, as it is illustrated in Fig 3-A.

**Fig 6.**
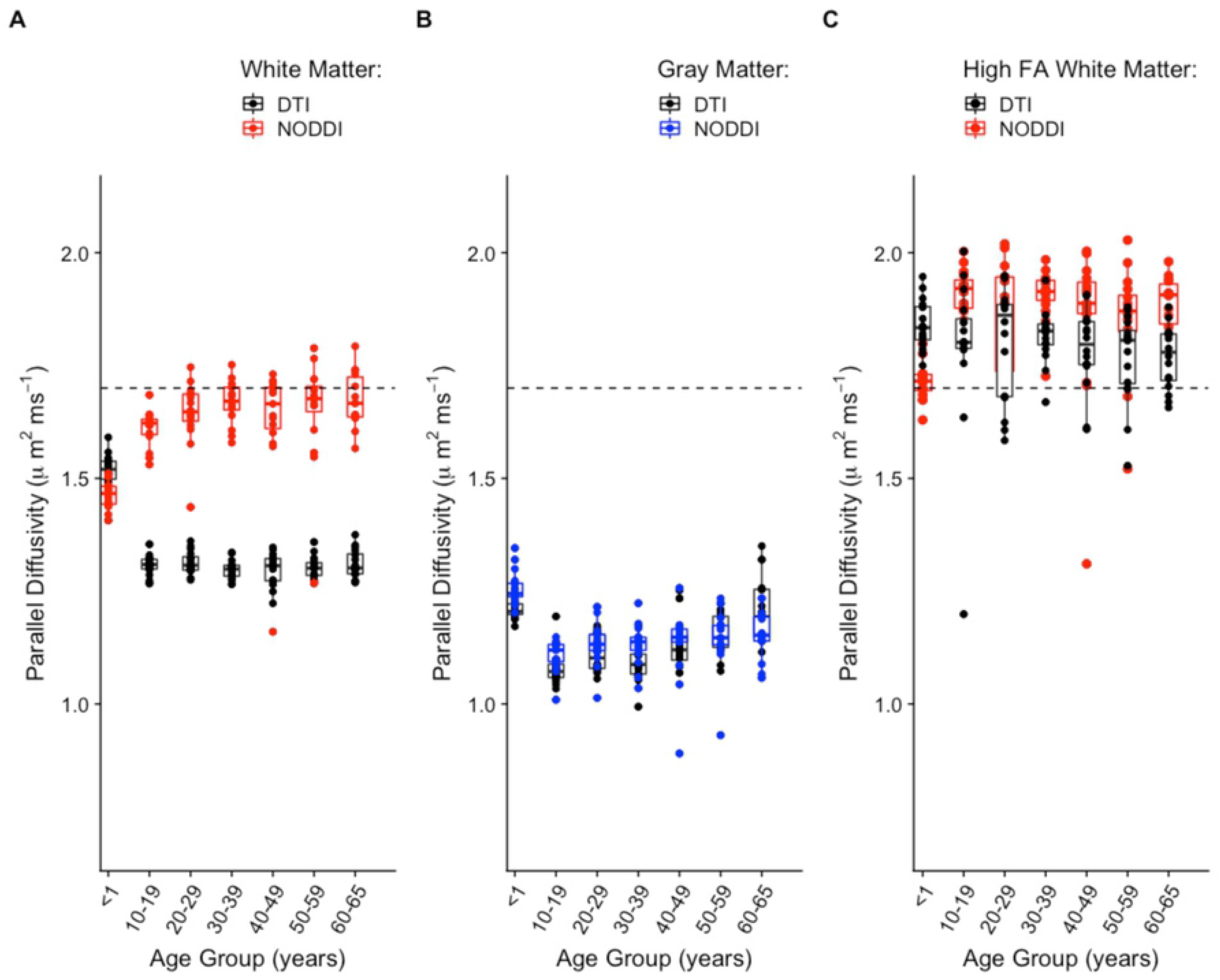
NODDI and DTI. Comparison of age trajectories between NODDI optimized parallel intrinsic diffusivity and DTI axial diffusivity in global white matter (**A**), global gray matter (**B**), and high FA white matter (**C**). The dashed horizontal line marks the NODDI default *d*_‖_ value.

## Limitations

### Assumed equal intra- and extra-cellular *d*_‖_

As mentioned in the introduction, another important assumption of the model is that of equal *d*_‖_ in the intra- and extra-cellular compartments. Thus, one of the limitations of this work is that it was carried out while maintaining this and other assumptions of the model.

In order to glimpse at the appropriateness of this assumption as it pertains to this work, a similar model residual optimization was done for the case where the extra-to intra-cellular parallel diffusivity ratio took on values different than 1. In this case, the model was adjusted so that the extra-cellular diffusivity was expressed as a fraction of the intra-cellular diffusivity value. The ratios ranged from 0.1 to 1.3 in 0.1 increments. In this case the number of fits increases dramatically for each subject (26×13=338), as do the memory and time requirements. Therefore, the analysis was restricted to two subjects, one infant and one adult, and for a single axial slice. Additionally, in order to circumvent the long fitting times using the Matlab tool box, for this part of the analysis the AMICO NODDI toolbox [38] was used instead.

Model RMS residuals were calculated for each of the 26 intra-cellular *d*_‖_ values in [0.5 *µ*m^2^⋅ms^−1^, 3.0 *µ*m^2^⋅ms^−1^] and each of the 13 extra-to intra-cellular *d*_‖_ ratio values in [0.1,1.3]. Average RMS residuals over WM and GM were plotted with respect to both, the intra-cellular *d*_‖_ and the ratio of extra-to intra-cellular *d*_‖_. These results are shown by the contour plots in Fig 7. Both in white and in gray matter, the regions of minimum residual values extend over several values in the two dimensions of the graphs. These poorly defined minima point to a multiplicity of solutions when constraints on the model diffusivities are not imposed. Similar results have been presented by other reports [23, 30], which show that unconstrained multi-compartment biophysical models lead to issues in parameter estimation. Particularly, the shape of the lowest residual regions in these contour plots is evocative the pipe-like structures for the fitting cost function landscapes of non-constrained multi-compartment models reported in [30] and [23].

**Fig 7.**
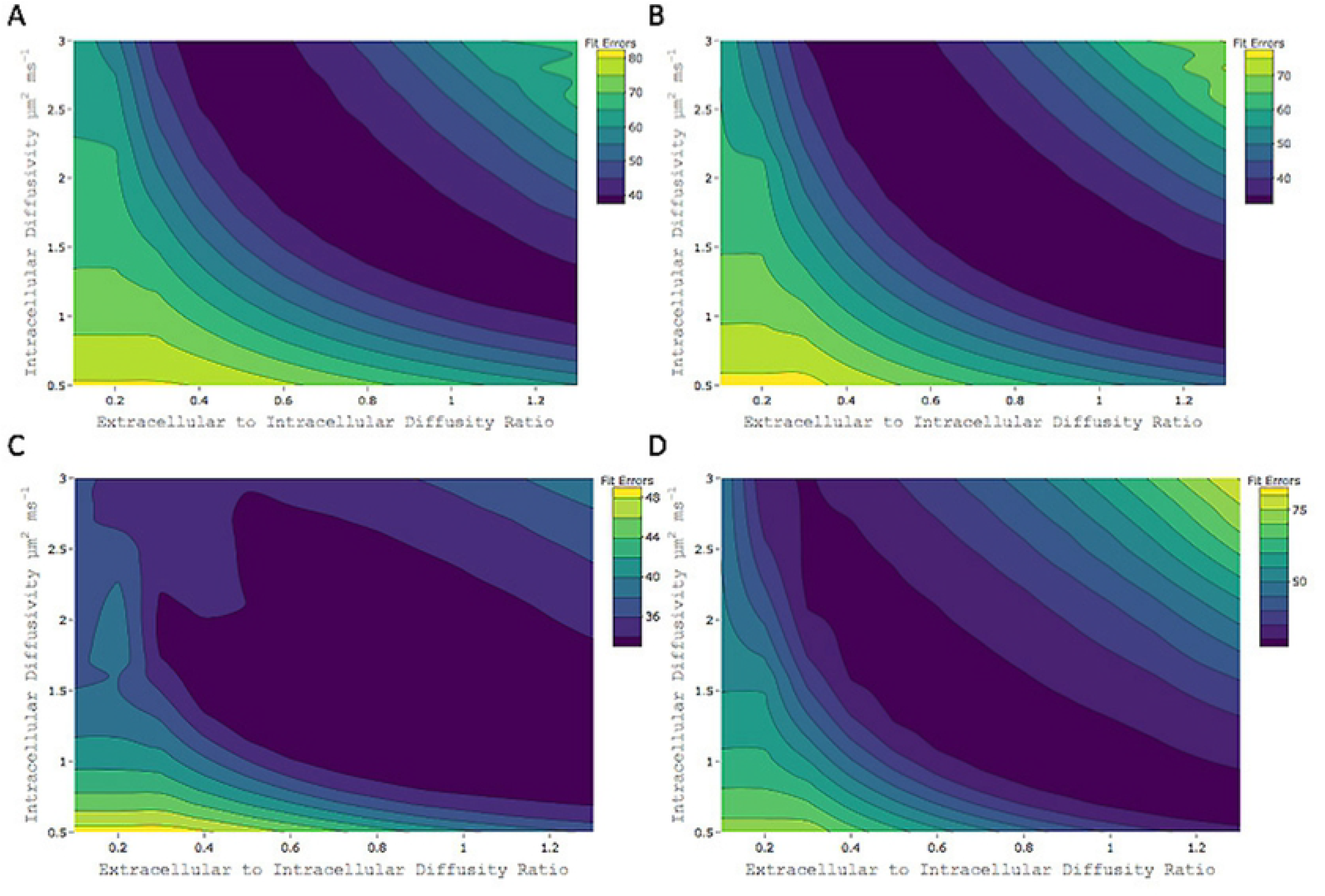
Model residuals and non-equal diffusivities. Fit errors (RMS residuals) of NODDI model with respect to both variation in intra-cellular *d*_‖_ and variation in the ratio of extra-to intra-cellular *d*_‖_. (**A**) Infant subject average fit errors over white matter. (**B**) Infant subject average fit errors over gray matter. (**C**) Adult subject average fit errors over white matter. (**D**) Adult subject average fit errors over gray matter.

### Generalizabily

Great effort was made in order to make this as an exhaustive analysis as possible in terms of the diversity of the data that was used. Yet, we acknowledge it is not fully generalizable to the wider scope of neuroimaging biophysical modeling diffusion research, for which it should consider, among others, conditions of pathology and ex-vivo experiments. Nonetheless, we believe that these results are highly informative considering the broad range of ages and imaging protocols investigated. Finally, this analysis was performed for Watson-NODDI only, not for other flavors of the technique which include Bingham-NODDI [39] or NODDIx [40].

## Conclusion

In this work, dependence of the estimated NODDI parameters on the parallel intrinsic diffusivity *d*_‖_ was observed. Optimal *d*_‖_ in white matter of the adult brain is similar to the currently used value but significantly lower in gray matter. Optimal *d*_‖_ is also lower than the default value for the newborn brain in white and gray matter. Effects of imaging protocol on the optimal *d*_‖_ were also observed. Caution should be used and optimization considered specially when conducting NODDI implementation in studies of gray matter and young populations (e.g. ¡10 years of age). It is important to consider that, despite its limitations, recent analysis suggests that NODDI metrics provide information that is congruent with histologically equivalent metrics [41].

## Acknowledgements

This study was supported in part by a core grant to the Waisman Center from the National Institute of Child Health and Human Development (IDDRC U54 HD090256). Additional support was provided by NIH grant R01 NS092870 (to AA), BRAIN Initiative R01-EB022883 (to NA), UW CPCP AI117924 (to NA), NIH grant R01AG037639, NIH grant R01AG027161, and NIH grant P50AG033514. JMG is supported by a UW-Madison SciMed Graduate Research Scholars Advanced Opportunity Fellowship, and the NSF Graduate Research Fellowship Program. Data collection for this study was supported by the Bill and Melinda Gates Foundation, grants P50 MH084051, R01 MH59785, the National Institute on Aging grants P01-AG020166 and U19 AG051426, and NIH grants P50 NIMH100031, R01 MH101504, R01 AG037639.

1 http://nitrc.org/projects/nodditoolbox

